# TMcin RiPP biosynthesis in the cellular membrane

**DOI:** 10.64898/2026.05.31.729130

**Authors:** Fauzia H. Nur, Ama N. Antwi, Seth W. Dickey

## Abstract

Ribosomally synthesized and post-translationally modified peptides (RiPPs) are a large category of natural products and a promising source for new medicines and applications in biotechnology. Their extraordinary diversity stems from the sequences of ribosomally synthesized precursor peptides and the wide repertoire of biosynthetic proteins that post-translationally modify and process the peptides. However, known precursor peptides and biosynthetic events have been characterized as soluble and occurring within aqueous environments in which RiPPs encounter the membrane primarily for secretion via transporters. Here, we report that the cell membrane is the central setting for the biosynthesis of TMcins, a recently discovered RiPP class with antimicrobial activity that contains a transmembrane helix (TMH) in the precursor peptide and mature product. We show that the ribosomally synthesized precursor TmcA is integrated in the producing cell membrane. We then uncovered the biosynthetic roles of gene products encoded on the TMcin biosynthetic gene cluster by integrating structure-prediction with an inducible TMcin biosynthesis platform. All TMcin post-translational modifications occurred in the membrane, for which three of four events were performed by intramembranous biosynthetic proteins. Finally, we assign an escort function to a previously uncharacterized membrane protein and provide insights into its evolution. Thus, our characterization of TMcin expands the setting for RiPP biosynthesis and provides a model for the biosynthesis of membrane-localized peptide natural products.

**Significance Statement:** Cells modify peptides to make natural products called RiPPs that exhibit a wide range of activities. Many RiPPs are antimicrobial and are important sources for discovering new drugs. Known RiPPs are produced by biosynthetic enzymes in aqueous environments. We characterized the recently discovered TMcin class of RiPPs that possess a transmembrane helix and are found across Gram-positive bacteria. Using a *Staphylococcus aureus* strain that naturally produces TMcin, we show that TMcin biosynthesis occurs in the membrane and elucidate the steps of TMcin production. This work establishes a model for understanding RiPP biosynthesis within a lipid environment that broadens our understanding of RiPPs, enables discoveries of new RiPPs, and provides a basis for the rational engineering of TMcins.

## Introduction

Microbes produce a vast array of natural products that continue to serve as a rich source for therapeutics and applications (*1*). Many are bacteriocins that kill or antagonize the growth of competing bacteria and offer novel treatments for drug-resistant infections (*2*). One large group of natural products are the ribosomally synthesized and post-translationally modified peptides (RiPPs) (*3–5*). These peptides are produced via ribosomal translation of a precursor peptide that is modified by biosynthetic proteins and enzymes, commonly encoded in close proximity within a biosynthetic gene cluster (BGC). The collective action of these proteins on a precursor peptide ultimately leads to the secretion of a mature and active RiPP. While the suite of biosynthetic proteins that process RiPPs perform a diverse range of chemical modifications, none have been characterized to act on transmembrane proteins (*3*, *4*).

Previously, we discovered the TMcin family of RiPPs (*6*). Strikingly, TMcins contain a transmembrane helix (TMH) in the mature and secreted antimicrobial product. After partitioning into a target cell membrane, TMcins oligomerize to form structured β-barrel pores that are surrounded by the TMHs, anchoring the pore in the target cell membrane. We also identified the BGC responsible for producing TMcin, which we found present across Gram-positive bacteria. Importantly, the TMcin BGC did not resemble other RiPP BGCs, which prevented its identification earlier by genome mining approaches.

Given the inclusion of the TMH, we became intrigued whether TMcin represents an intramembrane localized biosynthetic pathway and set out to experimentally characterize it. We investigated the cellular localization of TMcin biosynthesis within the producer cell and engineered an inducible expression system that we combined with predictive structures to map the processing roles of TMcin to BGC components and provide a model for TMcin biosynthesis.

## Results

### TMcin biosynthesis occurs at the membrane

We studied the model TMcin, TMcin-G1905, produced by *Staphylococcus aureus* G1905 (*6*, *7*). The TMcin-G1905 BGC is present on the pCT545 plasmid and encodes for several membrane proteins, including the structural gene *tmcA* (**Fig. 1*A***). Like TMcin, the precursor protein TmcA possesses a single TMH. We thus hypothesized that TMcin-G1905 is a RiPP biosynthesized at the bacterial membrane of producing cells. We applied our subcellular fractionation protocol, which we used previously to detect TMcin-G1905 in target cell membranes, to *S. aureus* G1905 and detected TMcin-G1905 in membrane preparations (**Fig. 1*B***). Interestingly, we also detected a second, higher molecular-weight band reactive to αTMcin-G1905 antisera specific to the TMcin C-terminal loop. Noting that this band was absent in the G1905 pCT545::pCURE strain lacking the TMcin BGC, we suspected that it represented the full-length TmcA precursor protein integrated in the membrane. We therefore fused the HA epitope tag to the N-terminus of TmcA and repeated the fractionation. Western blotting for the HA fusion and the TMcin-G1905 C-terminus confirmed the identity of the second band to be full-length TmcA (**Fig. 1*C***). Moreover and as reported earlier (*6*), only proteolytically processed TMcin-G1905 was present in secreted fractions (**Fig. 1*B***). Taken together, these data support the notion that TMcin-G1905 is processed at the membrane after ribosomal translation of the precursor protein TmcA, and that after maturation, TMcin-G1905 is secreted to diffuse away from the producing cell and reach the membranes of target cells.

**Fig. 1:**
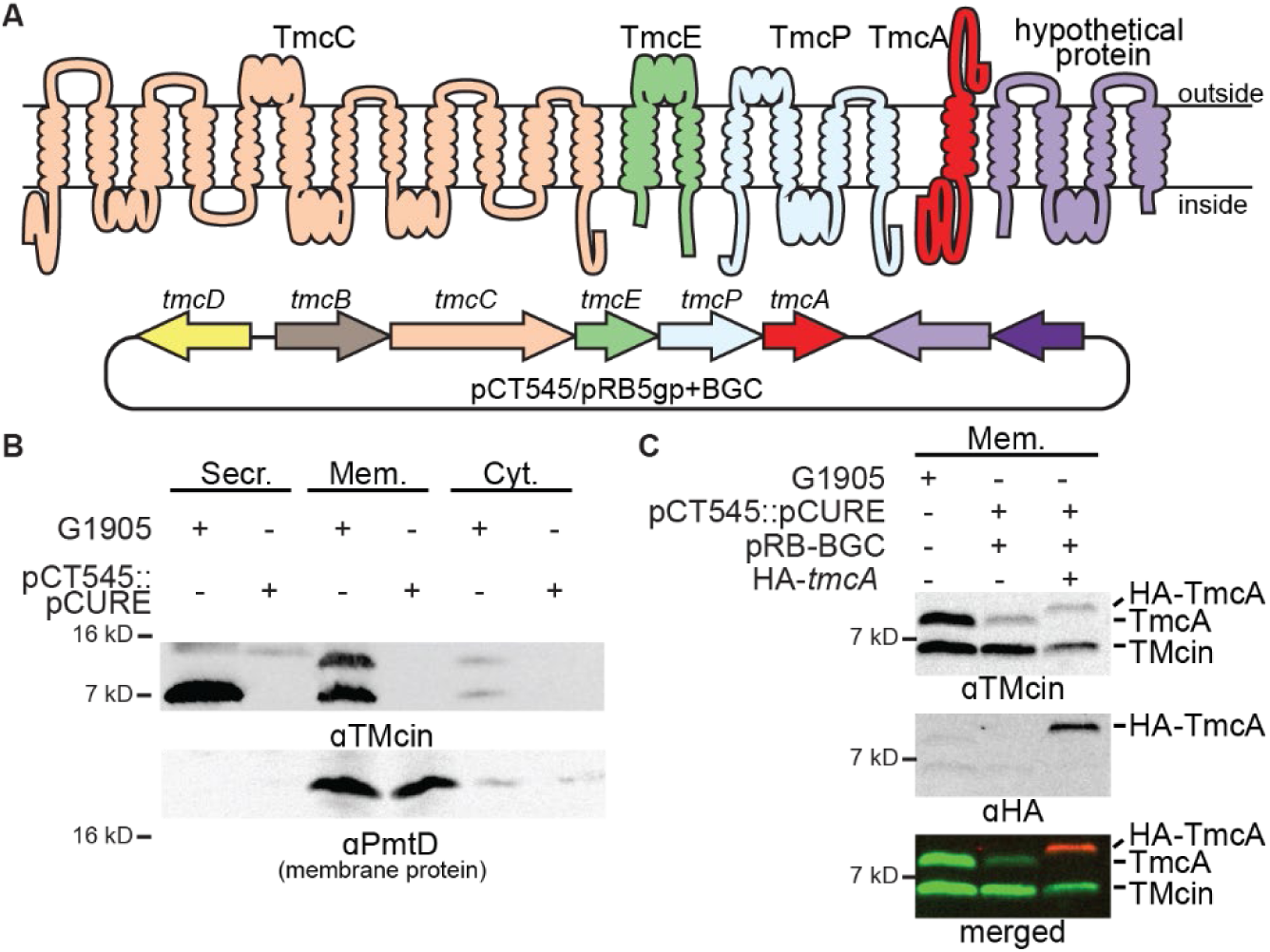
TMcin biosynthesis occurs in the cellular membrane. (*A*) Membrane topology diagram of TMcin biosynthetic proteins predicted to have transmembrane helices (above) encoded within the staphylococcal TMcin-G1905 BGC on pCT545 or the complementation plasmid pRB5gp+BGC (below). The function of open-reading frames with labeled gene names, except for *tmcA*, are described in this study. (*B*) Subcellular fractionation of *S. aureus* G1905 and cells cured of the pCT545 plasmid showing localization of TMcin and TmcA in cell membranes. The membrane protein PmtD (*39*) was used as a control for membrane fraction. (*C*) Western blots of cells expressing an HA-epitope fused to the N-terminus of TmcA, confirming that the higher molecular weight band in membrane preparations is TmcA. Secr., secreted fraction; Mem., membrane fraction; and Cyt., cytoplasmic fraction.

### Inducible expression of TMcin biosynthesis

Our prior characterization of the TMcin-G1905 bacteriocin entailed three post-translational processing events: site-specific proteolysis for leader peptide removal, disulfide bond formation between two universally conserved cysteine residues, and secretion into the extracellular space. To define the roles that the putative biosynthetic genes play in these events, we sought to engineer loss of function mutations within individual genes of the TMcin BGC and to characterize the TMcin-G1905 intermediates that accumulated as a result. We first attempted but failed to introduce loss of function mutations to the pRB+BGC plasmid we previously constructed that rescued TMcin-G1905 production in the G1905 pCT545::pCURE strain (*6*). We then reduced the overall plasmid size to improve transformation efficiency by deleting Gram-negative replication and antibiotic resistance genes (pRB5gp-BGC; gp for Gram-positive), however, we still failed to introduce loss of function mutations. We next adopted an inducible, two-plasmid strategy to mitigate the toxicity of TMcin-G1905 or its precursors. We placed the core operon that is conserved across TMcin BGCs and includes the structural *tmcA* gene behind a P_tet_ promoter (pKT17 BGC-core) to enable anhydrotetracycline (aTc)-inducible control, and we deleted that same operon from pRB5gp-BGC (pRB5gp BGC-Δcore, **Fig. 2*A***).

**Fig. 2:**
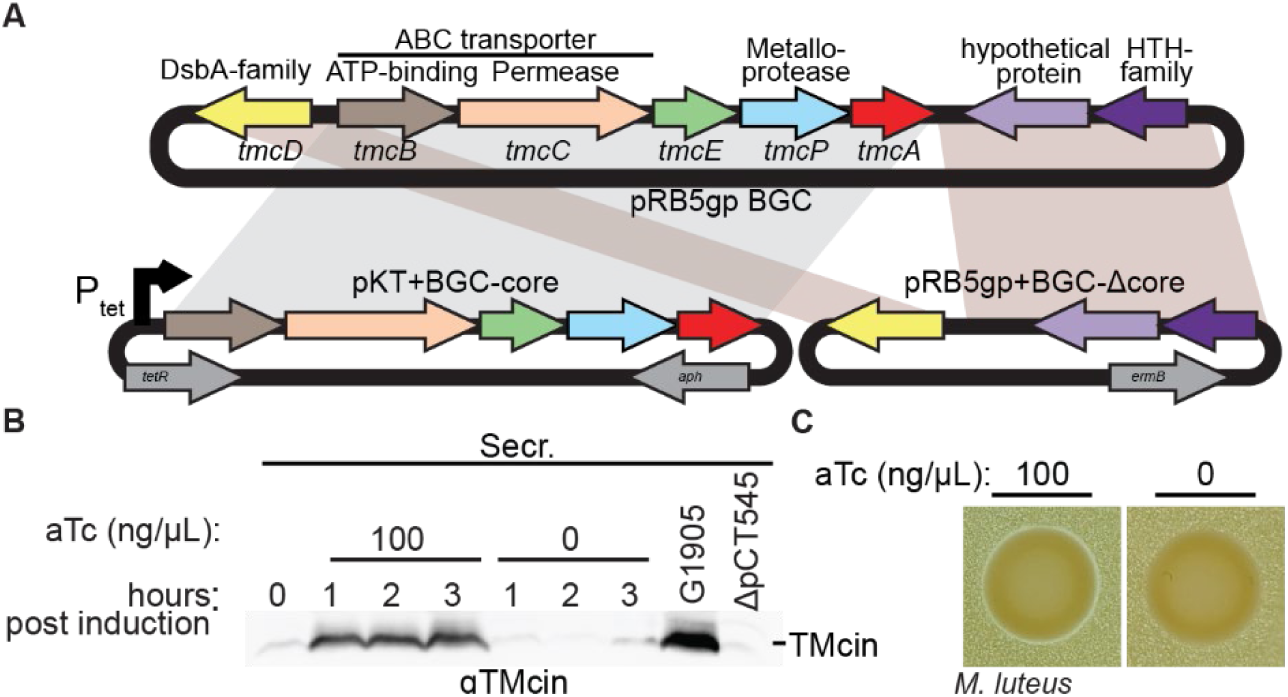
Inducible control of TMcin production. (*A*) G1905 P_tet_-TMcinBiosynthesis with a two-plasmid system in which the core operon of the TMcin-G1905 BGC encoding *tmcB*-*tmcA* is under control of a P_tet_ promoter on the engineered pKT plasmid. The remaining BGC genes, *tmcD* and the putative bicistronic regulatory operon (purple) are encoded on a second plasmid with native upstream sequences. (*B*) αTMcin western blots of G1905 P_tet_-TMcinBiosynthesis culture supernatants after inducing expression of the core operon by adding anhydrotetracycline (aTc). (*C*) Antimicrobial indicator plates containing *Micrococcus luteus* with or without aTc embedded in the agar. G1905 P_tet_-TMcinBiosynthesis cells were spotted on the plates. After incubation a zone of *M. luteus* growth clearance was present around the spot dependent on aTc present in the agar.

To allow for selection of both plasmids as well as induction using aTc, we generated a plasmid-free G1905 strain cured of pCT545 and deleted the chromosomal *tetM* and *tetL* genes (G1905 ΔpCT545 Δ*tetML*, see Methods). Finally, we transformed G1905 ΔpCT545 Δ*tetML* cells with both pRB5gp BGCΔcore and pKT17 BGC-core (G1905 P_tet_-TMcinBiosynthesis) and succeeded in detecting secreted TMcin-G1905 only after adding aTc (**Fig. 2*B***). In addition, inducing TMcin-G1905 biosynthesis conferred antimicrobial activity (**Fig. 2*C***), confirming inducible control of TMcin-G1905 biosynthesis. Using this system we were able to engineer precise loss-of function mutations throughout the TMcin BGC (see below).

### Leader peptide removal

*S. aureus* G1905 TmcA includes a 45-residue leader peptide that is removed during maturation, consistent with RiPPs more generally (*8*). We previously proposed that the fourth open reading frame (ORF), which we termed *tmcP*, in the core operon encodes the site-specific protease because it included a HExxH zincin metalloprotease motif (*9*, *10*). Using hmmscan (*11*), we found that TmcP is a member of PF11667 family. This family is defined by proteins with the domain of unknown function 3267 (DUF3267; **Fig. 3*A***, **Table S1**) that was proposed to achieve a zincin-like metalloprotease fold (*12*). To provide additional insights, we generated a predicted structural model using AlphaFold3 (AF3) (**Fig. 3*B***, **Fig. S2*A***) and used it to search for structural homologs (*13*). This resulted in one hit (pdb 3B4R) in the PDB100 database to the *Methanocaldococcus jannaschii* Site-2 protease (mjS2P) (*14*) (**Fig. 3*B***). Corroborating this, the active site residue architecture of TmcP is well preserved, including the distal Asp and Pro, which are hallmarks of the S2P-zincin clan (MEROPS MM) (*10*) and the DUF3267 HMM profile (**Fig. 3*A***). Interestingly, TmcP lacks the last two TMHs 5 and 6 of mjS2P that are proposed to serve as a substrate gate regulating access to the proteolytic active site (**Fig. 3*B***). Moreover, TmcP had a TM score > 0.5, indicating a similar fold (*15*), to >97% of DUF3267 pre-generated AF predictions, and 88% of DUF3267-containing proteins are predicted to have four TMHs (**Table 1**). Using our TMcin biosynthesis assay, we tested whether the TmcP catalytic motifs were necessary for TmcA cleavage by mutating the catalytic histidine codon (H^68^) of the H^68^EYMH motif and the distal aspartate codon (D^158^) with alanine codons (*tmcP* AA). This abrogated production of TMcin-G1905 and led to the accumulation of full-length TmcA in cell membranes (**Fig. 3*C*, Fig. S1**). Thus, TmcP is responsible for site-specific removal of the TmcA leader peptide, is a member of the S2P clan of intramembrane proteases, and is representative of the DUF3267-containing PF11667 family within the S2P clan.

**Fig. 3:**
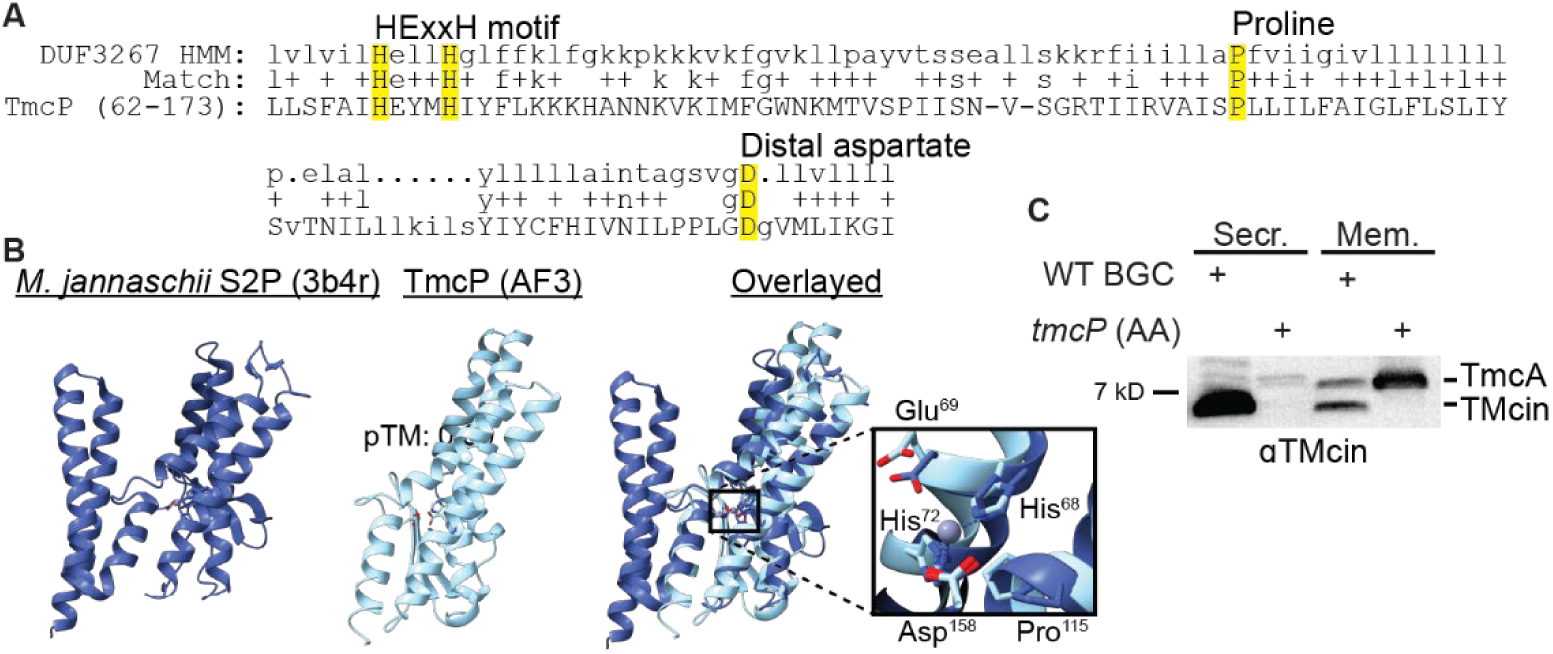
TmcP is a site-specific member of the S2P-family responsible for TMcin leader peptide removal. (*A*) Alignment of TmcP subsequence with the DUF3267 HMM profile. HMM residues with >50% emission probability are highlighted in yellow. (*B*) Cartoon drawing of the mjS2P crystal structure (3b4r, left), an AF3-generated structure of TmcP (middle, pTM 0.89), and with both structures overlaid (right). The inset magnifies the catalytic site and Zn^2+^ ion coordination. (*C*) Subcellular fractionation and western blot of TMcin after induction of TMcin biosynthesis in which the catalytic histidine codon (H^68^) and the distal aspartate codon (D^158^) of *tmcP* were mutated to alanine.

**Table 1.**
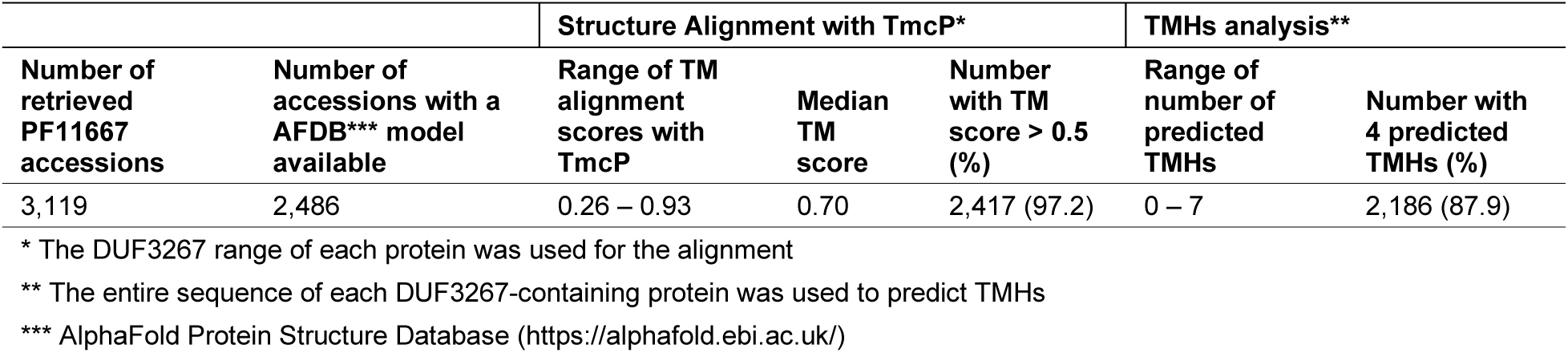
Comparison of PF11667/DUF3267-containing proteins to TmcP.

| Number of retrieved PF11667 accessions | Number of accessions with a AFDB*** model available | Structure Alignment with TmcP* |  |  | TMHs analysis** |  |
| --- | --- | --- | --- | --- | --- | --- |
|  |  | Range of TM alignment scores with TmcP | Median TM score | Number with TM score > 0.5 (%) | Range of number of predicted TMHs | Number with 4 predicted TMHs (%) |
| 3,119 | 2,486 | 0.26 – 0.93 | 0.70 | 2,417 (97.2) | 0 – 7 | 2,186 (87.9) |

\* The DUF3267 range of each protein was used for the alignment
\*\* The entire sequence of each DUF3267-containing protein was used to predict TMHs
\*\*\* AlphaFold Protein Structure Database (https://alphafold.ebi.ac.uk/)

### Disulfide-bond formation

TMcin has one intramolecular disulfide bond between two universally conserved cysteine residues that loops the C-terminal end to the top of the TMcin TMH (*6*). We previously reported that treating TMcin with reducing reagents led to a gel-migration shift, which we successfully replicated here (**Fig. 4*A***), and we interpreted that this shift reflected breaking the disulfide bond. To test this interpretation more rigorously, we used a sulfhydryl-reactive maleimide with a discrete poly-ethylene glycol chain length (m-dPEG_36_-MAL) that specifically labels the free thiols of reduced cysteine residues to further slow the gel-migration of reduced TMcin. Purified TMcin-G1905 failed to interact with m-dPEG_36_-MAL, consistent with an intact disulfide bond. However, TMcin-G1905 pre-treated with the reducing reagent TCEP resulted in an additional discrete and slower-migrating band on SDS-PAGE (**Fig. 4*A***). Inducing expression of G1905 P_tet_-TMcinBiosynthesis cells carrying mutations of the *tmcA* cysteine codons (C25 and C50) to alanine (*tmcA* AA), resulted in a similar gel-migration shift as reduced TMcin-G1905 (**Fig. 4*B***). Thus we conclude that the gel-migration shift faithfully reports on the status of the TMcin disulfide bond. Importantly, we also observed a gel-migration shift for both TMcin and the precursor TmcA in cell membrane fractions as a result of the *tmcA* AA mutation (**Fig. 4*B***), indicating that disulfide bond formation occurs within the membrane.

**Fig. 4:**
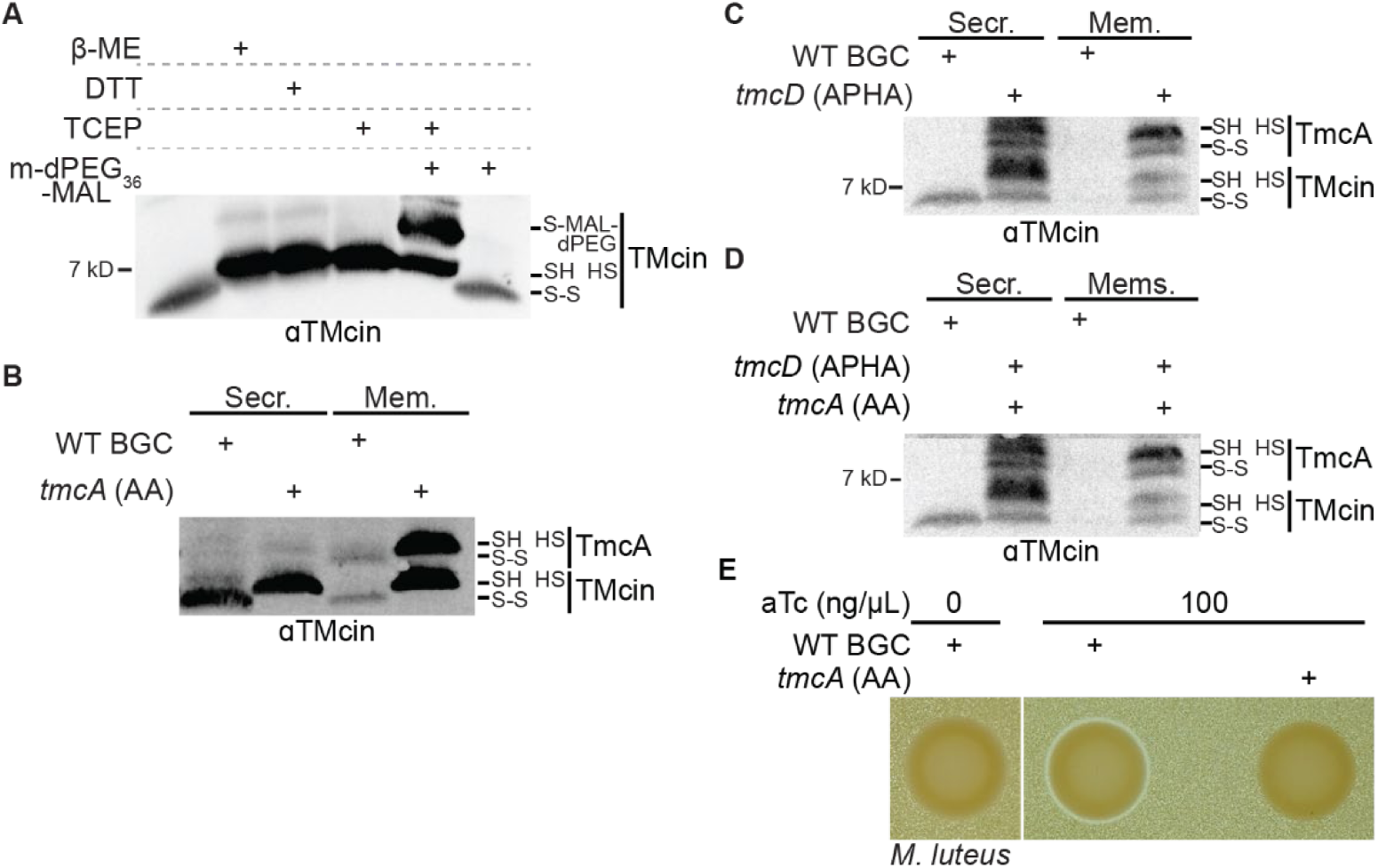
The DsbA-like protein TmcD forms the TMcin disulfide bond. (*A*) Western blot of purified TMcin-G1905 treated with reducing reagents and/or the thiol specific maleimide conjugated to PEG with a discrete chain length. (*B-D*) Subcellular fractionation and western blot of TMcin after induction of biosynthesis in which the (*B*) *tmcA* cysteine codons were mutated to alanine, (*C*) the catalytic cysteine codons of *tmcD* were mutated to alanine, or (*D*) both. (*A-D*) All samples were separated using urea-polyacrylamide gels. (*E*) Antimicrobial indicator assays showing that disruption of the TMcin disulfide bond results in a loss of antimicrobial activity.

Staphylococcal TMcin BGCs encode a DsbA-family protein oxidoreductase, *tmcD*, that includes a CXXC motif common in oxidoreductase enzymes. This motif is identical to that of *E. coli* DsbA (CPHC), in which the His residue reduced the pKa of the cysteine thiols, increasing the oxidizing potential of DsbA to form disulfide bonds within other proteins (*16*). We mutated the *tmcD* catalytic cysteine codons with alanine codons (*tmcD* APHA) and this led to a conversion of TMcin-G1905 from a single band to several bands (**Fig. 4*C***). The band with the greatest intensity corresponded to reduced TMcin, however, we also detected a faster migrating band that could be due to spontaneous formation of disulfide bonds and slower migrating bands (**Fig. S1**) that may represent other cysteine oxidation byproducts or conjugates. To test that these other populations depended on the TMcin cysteine residues, we combined the *tmcA* AA and *tmcD* APHA mutations. This resulted in two bands that corresponded to reduced TMcin and TmcA (**Fig. 4D**). Finally, although secreted TMcin levels were similar, the G1905 P_tet_-TMcinBiosynthesis cells with the *tmcA* AA mutation failed to clear *M. luteus* on antimicrobial indicator plates, demonstrating that the TMcin disulfide bond is required for antimicrobial activity and is a critical step in the maturation of TMcin (**Fig. 4*E***).

### Secretion

The *tmcB* and *tmcC* genes encode for ATP binding and permease domains, respectively, that together form a putative ABC transporter. The AF3 predicted structure includes a dimer of TmcB subunits in complex with one copy of the 12-TMH TmcC (**Fig. 5*A*, Fig. S2*B***). TmcC adopted an apparent single-chain type V permease fold (*17*, *18*) in which the most of the transporter would be embedded in the membrane. Because a single TMH is sufficient to anchor proteins to the membrane, we hypothesized that TmcBC extracts TMcin-G1905 from the membrane for secretion, analogous to the extraction of sterols by the mammalian ABCG5/8 transporter (*19*) or of amphipathic peptides by the *S. aureus* PmtCD transporter (*20*, *21*). We mutated the *tmcB* catalytic glutamate codon after the Walker B motif to a glutamine codon (*tmcB* E156Q) to disrupt ATP hydrolysis. Consistent with a secretory role, the *tmcB* mutation led to a loss of TMcin-G1905 in the secreted fraction even though the levels of TMcin-G1905 and TmcA in the membrane fraction remained similar (**Fig. 5*B***).

**Fig. 5:**
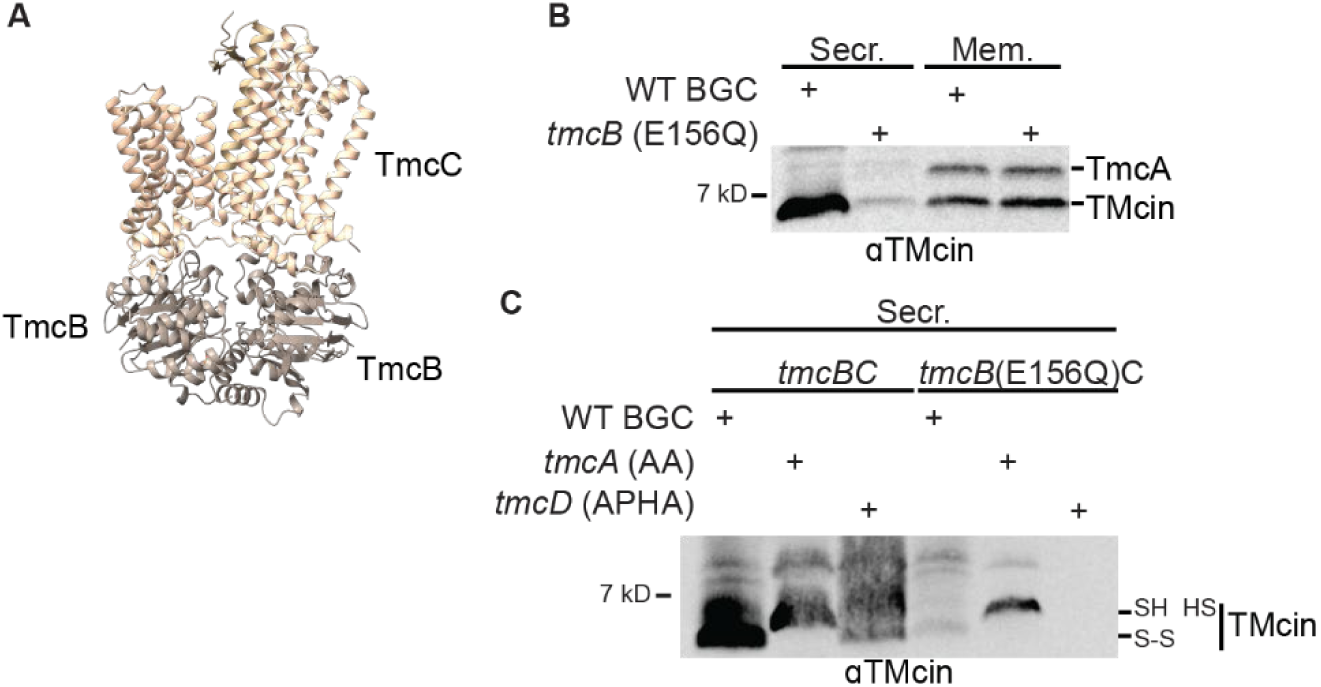
The TmcBC ABC transporter is required for TMcin secretion. (*A*) Cartoon drawing of a AF3-generated model of the TmcBC transporter with two copies of TmcB (brown) and one copy of TmcC (tan, pTM 0.81, piTM 0.77). (*B*) Subcellular fractionation and western blot image of TMcin after induction of biosynthesis in which the glutamate codon of the Walker B motif in *tmcB* was mutated to glutamine or (*C*) secreted fractions separated using a urea-polyacrylamide gel with mutations interfering with disulfide bond formation combined with the *tmcB* catalytic mutations.

Despite disrupting disulfide bond formation, TMcin was still present in culture filtrates (**Fig. 4*C***). We next asked whether this depended on export by TmcBC. Combining the *tmcB* (E156Q) catalytic mutation with either the *tmcA* (CC-AA) or the *tmcD* (APHA) mutations led to a reduction of TMcin in culture filtrates (**Fig. 5*C***), demonstrating that TmcBC can transport TMcin regardless of disulfide bond formation.

### Escort

The remaining ORF, which we term *tmcE*, of the BGC core operon encodes for a protein with two membrane spanning domains but with no other identifiable motifs or homology to characterized protein families. Nonetheless, *tmcE* is necessary for TMcin biosynthesis. We disrupted *tmcE* by replacing serine 10 and leucine 11 codons with stop codons (*tmcE* 2xstop). This resulted in a phenocopy of the *tmcP* AA mutation where we only observed the full-length TmcA intermediate in the cell membrane fraction, indicating that TmcE promotes proteolysis by TmcP (**Fig. 6*A***). Supporting this notion, we generated a confident predictive structural model of TmcE in complex with TmcA and TmcP (**Fig. 6*B*, Fig. S2*D***). Intriguingly, TmcE is in a similar position as the missing gating helices. In fact, TmcE showed structural similarity to a subset of DUF3267 proteins: 381 (15.3 %) of DUF3267 pre-generated AF models yielded a TM-score >0.5 when aligned to the TmcE AF3 predicted structure and 22 (0.9 %) had TM-scores >0.7 (**Fig. 6*C***, **Table 2**). In addition, DUF3267-proteins with 6 predicted transmembrane helices are enriched for TM scores >0.5 or >0.7 (**Table 3**).

**Fig. 6:**
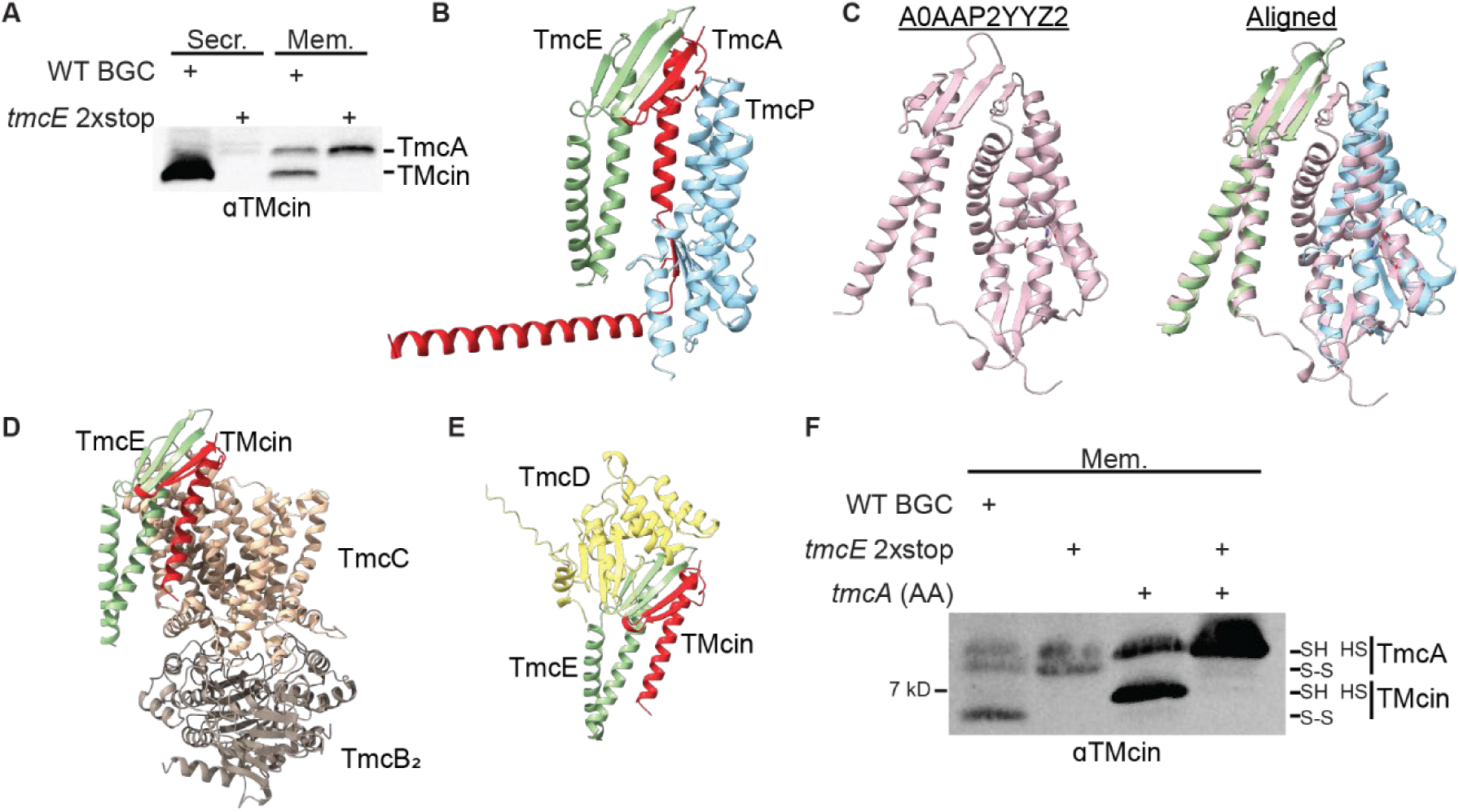
TmcE is an escort protein of TmcA and TMcin and is required for leader peptide removal. (*A*) Subcellular fractionation and western blot image of TMcin after induction of biosynthesis in which the *tmcE* ORF was disrupted with two premature stop codons. (*B*) Cartoon drawing of the TmcA, TmcE, and TmcP AF3-generated complex structural model (pTM 0.83, piTM 0.87), (*C*) Structural alignment of TmcE with the top-scoring DUF3267-containing protein (UniProt #A0AAP2YYZ2). The overlay with TmcP is also shown (TM 0.73 and 0.53 for TmcE and TmcP as references, respectively). (*D*) Cartoon drawing of AF3-generated complex structural models of TMcin, TmcE, and TmcBC (pTM 0.74, piTM 0.70) and, (*E*) TMcin, TmcE, and TmcD (pTM 0.80, piTM 0.76). (*F*) Western blot of TMcin in the membrane fraction after induction of biosynthesis in which the *tmcE* ORF was disrupted. Samples were using a urea-polyacrylamide gel.

**Table 2.** Comparison of PF11667/DUF3267-containing proteins to TmcE.

| Structure Alignment with TmcE* |  |  | Analysis of TMHs |  |  |
| --- | --- | --- | --- | --- | --- |
| Range of TM alignment scores with TmcE | Median TM score | Number with TM score > 0.5 (%) | Number with 6 predicted TMHs (%) | Number with 6 predicted TMHs & TM score > 0.5 (%) | Number with 6 predicted TMHs, TM score > 0.5, & with TMHs 5 & 6 that aligned to TmcE |
| 0.29 – 0.73 | 0.43 | 381 (15.3) | 141 | 131 (92.9) | 131 |

\*The entire sequence of each DUF3267-containing protein was used for alignment with TmcE

**Table 3.** Contingency tables of PF11667/DUF3267-containing proteins alignment to TmcE.

TM > 0.5
| DUF3267 proteins | TM score > 0.5 | TM score ≤ 0.5 | Total |
| --- | --- | --- | --- |
| With 6 TMHs | 131 | 10 | **141** |
| Not with 6 TMHs | 250 | 2,095 | **2,345** |
| **Total** | **381** | **2,105** | **2,486** |

Fisher's exact test p value < $2.2 \times 10^{-16}$ TM > 0.7
| DUF3267 proteins | TM score > 0.7 | TM score ≤ 0.7 | Total |
| --- | --- | --- | --- |
| With 6 TMHs | 21 | 120 | **141** |
| Not with 6 TMHs | 1 | 2,344 | **2,345** |
| **Total** | **22** | **2,464** | **2,486** |

Fisher's exact test p value < $2.2 \times 10^{-16}$

We also generated a complex between TmcE, TMcin, and TmcBC (**Fig. 6*D***, **Fig. S2*E***) as well as TmcE, TMcin, and TmcD (**Fig. 6*E*, Fig. S2*F***). However for the latter, the TMcin cysteine residues were not located near the TmcD catalytic residues and *tmcE* was not required for disulfide bond formation (**Fig. 6*F***). This suggests that TmcE remains in complex with TMcin after proteolysis and is also required for secretion. We therefore propose that TmcE performs an “escort” role in which TmcE interacts with TMcin intermediates within the membrane to promote leader peptide removal and secretion.

## Discussion

RiPPs have emerged as a source of structurally and functionally diverse natural products, including antimicrobials (*3*, *4*). This is underpinned by a diverse set of biosynthetic proteins and RiPP pathways. Here we add to our understanding of RiPP biosynthesis by describing TMcin biogenesis. TMcin is notable in that it possesses a TMH that allows it to form oligomeric structured pores in target cell membranes (*6*). Using TMcin-G1905 produced by *S. aureus* G1905 as a model, we find that the TMH is also critical in guiding TMcin biosynthesis by integrating TMcin and the precursor peptide TmcA into the cell membrane. We then engineered an inducible expression system that enabled systematic mutagenesis of the TMcin BGC, which we used to assign four key processing roles: 1) proteolysis by TmcP to remove the leader peptide, 2) disulfide bond formation, necessary for antimicrobial activity, by the DsbA-like protein TmcD, 3) TMcin secretion by the ABC transporter TmcBC, and 4) escort by the previously uncharacterized protein TmcE, which we propose to interact with TmcA and TMcin throughout biosynthesis (**Fig. 7**).

**Fig. 7:**
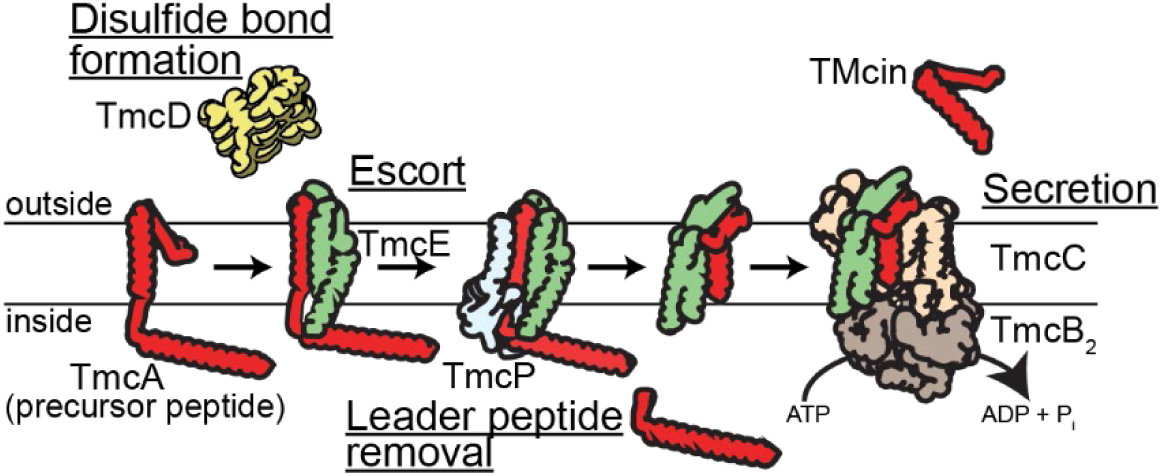
Model of TMcin biosynthesis in the cellular membrane. The precursor TmcA undergoes disulfide bond formation by TmcD, interacts with the escort protein TmcE, and is cleaved by TmcP to form TMcin. TmcE-TMcin interacts with transporter TmcBC, after which TmcE dissociates and TMcin is secreted.

Other RiPPs, although not embedded within the membrane, need to be transported across the membrane for secretion (*22*, *23*). In several cases, transport is coupled with additional processing events occurring near the membrane. For instance the PCAT transporters include a proteolytic domain that removes the leader peptide before secretion and the lanthipeptide nisin is modified by a protein complex that is anchored to the membrane by the NisT transporter (*24–28*). Nonetheless, TMcin biosynthesis is distinct in that the membrane is the central setting throughout biosynthesis. The TMH is processed at both ends (proteolysis and disulfide bond formation) and the TMH itself is extracted from the membrane for secretion.

Our characterizations also provide insights into the evolution of this membrane-embedded RiPP pathway. We discovered that TmcP is a S2P intramembrane protease (MEROPS clan MM) with the DUF3267. It, and most other DUF3267 proteins, possess only 4 TMHs, conspicuously lacking the two C-terminal TMHs 5 and 6 of the mjS2P and the *Bacillus subtilis* S2P, SpoIVB (*14*, *29*). These helices are proposed to undergo large movements, serving as a gate for transmembrane substrates. Interestingly we generated a ternary complex of TmcP, TmcA, and TmcE. TmcE is a 2-TMH membrane protein and occupied a similar position of the missing helices. Genetic ablation of *tmcE* led to accumulation of the precursor protein in cell membranes, phenocopying *tmcP* catalytic-dead mutations and demonstrating that TmcE controls TMcin proteolysis. We also found a subset of DUF3267 proteases, highly enriched for proteins with 6 TMHs, that structurally aligned with TmcE, including a three-stranded β-sheet between the two TMHs. Our results have two important implications: 1) regulatory functions analogous to TmcE may extend to other DUF3267 proteases with 4 TMHs and 2) while we cannot rule out convergent evolution, our results support a model in which TmcE and TmcP derived from a common 6-TMH DUF3267 ancestral protein.

Such an event would have allowed the TmcE ancestor to maintain an interaction with the proto-TMcin proteolytic product. Indeed, we also generated structural models of TmcE in complex with TMcin and TmcD as well as with TMcin and the TmcBC transporter. Thus it appears that TmcE plays a role throughout TMcin biosynthesis, which may have allowed TMcin to gain pore-formation functionality while preventing homo-oligomerization in the producing cell until after processing and delivery to a transporter for secretion.

Our findings also have implications for the preconditions of TMcin BGC mobilization on a plasmid in staphylococci. A disulfide bond between universally conserved cysteine residues (TMcin-G1905 C25 and C50) is required for antimicrobial activity (**Fig. 4*B*,*E***) and the TmcD DsbA-like gene product encoded by *tmcD* is responsible for promoting disulfide bond formation (**Fig. 4*C***). However, *tmcD* is missing in non-staphylococcal TMcin BGCs (*6*). This may reflect that DsbA enzymes are thought to be promiscuous and potentially allowing for expression of distally encoded oxidoreductases to achieve TMcin disulfide bond formation (*30*, *31*). Notably, *S. aureus* produces few proteins with disulfide bonds (*32*, *33*) and while *S. aureus* encodes a conserved DsbA protein on the chromosome, it is insufficient for robust TMcin disulfide bond formation (**Fig. 4*C***). Interestingly, the TmcD catalytic motifs are identical to those of *Escherichia coli* DsbA, which yield a stronger oxidative potential than those of *Sa*DsbA (*34*). Thus, it is plausible that staphylococci lack the capacity to form the TMcin disulfide bond, making the inclusion of *tmcD* in the BGC necessary for producing antimicrobially active TMcin in staphylococci. Alternatively, given that reduced TMcin can be secreted, other organisms may secrete antimicrobially inactive TMcins, perhaps relying on environmental oxidation to form a disulfide bond.

This study did not include analysis of the putative regulatory operon nor an investigation into the mechanism of self-immunity. However, the TmcE escort function, along with a predicted interaction via an extended β-sheet with the C-terminal segment of TmcA or TMcin would compete for and prevent TMcin homo-oligomerization and, thereby, pore formation. We presume that the escort function of TmcE is key to preventing autotoxicity during TMcin biosynthesis.

In summary, we provide insights into how a RiPP with a TMH is processed and how biosynthesis evolved in the unique membrane environment. This expands our understanding of natural product biosynthesis and will enable heterologous systems to facilitate mechanistic studies and recombinant production platforms of the TMcin family.

## Methods

### Bacterial strains and growth conditions

*S. aureus* strains were grown at 37°C, shaking at 180 rpm in tryptic soy broth (TSB) or statically on tryptic soy agar. Erythromycin and Kanamycin were added as needed to maintain selection of plasmid at 10 and 100 µg/ml, respectively. Plasmids used in this study are listed in *SI Appendix*, Table 3.

### Strain engineering

The pCT545 plasmid was removed from *S. aureus* G1905 without the need of a heterologous plasmid to continuously provide the antitoxin of the pCT545 toxin/antitoxin (TA) system (G1905 ΔpCT545). The putative pCT545 TA system was amplified from G1905 genomic DNA with primers (**Table S3**) that substituted two stop codons for the toxin start codon and was inserted into the pIMAY EcoRV site (pIMAY CT545-AT). This plasmid was introduced into G1905 by electroporation and single colonies were cultured and screened by PCR for spontaneous loss of the pCT545 plasmid. The pIMAY CT545-AT was removed by shifting to the non-permissive growth temperature of 37 °C.

The *tetM* and *tetL* genes were deleted in G1905 ΔpCT545 by allelic exchange to allow for intracellular accumulation of aTc during induction. The *tetML* locus and flanking 1kb homology arms was amplified from G1905 genomic DNA (Table S2) and the PCR product was inserted into the EcoRV site of pIMAY. *tetM* and *tetL* were deleted by inverse PCR (pIMAY 1kb-Δ*tetML-*1kb)

### Molecular cloning and mutagenesis

Oligonucleotides and engineered plasmids used in this study are listed in (**Tables S2 and S3**). PCR products were amplified using Q5 polymerase (NEB). For mutagenesis by inverse PCR, the PCR product was circularized using T4 PNK and T4 ligase (NEB). Clones were confirmed by restriction digestion or PCR, and all sequences were validated by Sanger (Azenta) or nanopore (Plasmidsaurus) sequencing.

pKT17 was engineered from pKX17 via pKX15 (*35*). First, pKX17 was engineered to introduce a multiple cloning site similar to pTX17 (*21*) using pKX15 as a template for inverse PCR and recircularization by T4 PNK and T4 ligase (NEB). The *xylR* regulator and P_xylA_ promoter from pKX17 were removed by inverse PCR and the PCR product was digested with ApaI. The *tetR* regulator and P_tet_ promoter from pIMAY were amplified, digested with ApaI and treated with T4 PNK. The two PCR products were ligated and one NdeI restriction site in *tetR* was removed by inverse PCR to yield pKT17 for anhydrotetracycline (aTc) inducible expression.

pRB5gp+BGC was generated by inverse PCR using pRB573+BGC (*6*) as a template for inverse PCR to delete the Gram-negative origin of replication and the β-lactamase antibiotic resistance marker. pRB5gp-BGCΔcore was generated from pRB5gp+BGC by inverse PCR to delete *tmcB-tmcA.* The BGC core operon was amplified from G1905 genomic DNA and inserted into pKT17 at the NdeI and NotI sites.

### Induction of TMcin biosynthesis

Overnight starter cultures were diluted to an OD_600_ of 2.0 in TSB supplemented with erythromycin and kanamycin and were cultured at 37 °C, 180 rpm for 2 hours. aTc was added to a final concentration of 1 µg/ml, and the cultures were incubated for an additional 1.5 hours.

### Subcellular fractionation

Subcellular fractionation was performed similarly to before (*6*). Briefly, secreted fractions were obtained by centrifuging bacterial cultures at 15,000 rpm for 2 minutes, passing the supernatant through a 0.2 µm PES filter, and concentrating the filtrates 20-fold by TCA precipitation, washing twice with cold-acetone, and resuspending the pellet with 2X SDS-sample buffer. For cytoplasmic and membrane fractions, the cell pellets were resuspended in 1X TBS and lysed using acid-washed ≤100 µm glass beads (Sigma Aldrich #G4649). The beads were gently pelleted by centrifugation, and the supernatant was ultracentrifuged using a Beckmann Type 42.2 Ti fixed-angle rotor at 20,000 rpm for 15 minutes. The supernatant was collected as the cytoplasmic fraction and the pellet was resuspended with 2X SDS-sample buffer as the membrane fraction.

### Protein analyses

Protein samples were resolved by SDS-PAGE in 1X tricine running buffer using 16% polyacrylamide gels (*36*). To detect gel shifts of TMcin-G1905 based on the disulfide bond status, samples were prepared in SDS sample buffer without reducing reagents, unless noted, and were separated using 12% polyacrylamide gels with 6 M urea. For total protein staining, gels were stained using the Imperial Protein Stain (ThermoFisher #24615) following the manufacturer’s instructions. For western blotting, semidry transfer was used to blot proteins onto 0.2 µm nitrocellulose membranes in transfer buffer (50 mM Tris, 40 mM glycine pH 9.2, 20% methanol) at 25 V for 30 minutes. TMcin-G1905 was detected using a TMcin C-terminal–specific rabbit polyclonal antiserum (*6*) followed by goat-αrabbit-IR800CW IgG. The HA tag was detected using the rat monoclonal antibody 3F10 (Roche #11-867-423-001) followed by goat-αrat-IR680RD. Membranes were imaged using an Azure 600 imager (Azure Biosystems).

### Antimicrobial indicator plates

*S. aureus* G1905 P_tet_-TMcinBiosynthesis strains were cultured in TSB supplemented with erythromycin and kanamycin. The cells were washed three times with sterile TSB to remove residual antibiotics, and 10 µL was spotted on TSA indicator plates prepared with 5*10^6^ CFU/mL *Micrococcus luteus* with and without 0.1 μg/mL aTc embedded in the agar. The plates were incubated at 37°C overnight, and the presence of clear zones around the spots was assessed for antimicrobial activity.

### TMcin-G1905 purification

TMcin-G1905 was purified as before (*6*). *S. aureus* G1905 Δ*agr* cells were cultured shaking at 220 rpm in TSB at 37 °C. Cells were harvested by centrifugation (Eppendorf Centrifuge 5810 R), washed with deionized H₂O, and washed in 8 M urea. The 8 M urea wash was filtered through a 0.2 µm polyethersulfone (PES) filter.

TMcin-G1905 was purified from the wash-filtrate by reverse-phase chromatography using C18-solid phase extraction (SPE) columns (Waters #WAT036905) followed by size-exclusion chromatography as previously described. TMcin-G1905 preparations were lyophilized, resuspended to 20 µM with 50% acetonitrile and 0.1% trifluoroacetic acid, and stored at –80 °C.

### Disulfide bond analysis

Methoxy-dPEG36-maleimide (Vector Laboratories, #QBD-10931), a discrete-length PEG bearing a sulfhydryl-reactive maleimide, was used to label reduced TMcin-G1905. For the PEGylation reactions, 4 µM purified TMcin-G1905 was pretreated with or without 10 mM tris(2-carboxyethyl) phosphine (TCEP) in 4X SDS sample buffer and heated to 95 °C for 5 minutes. The reactions were supplemented with 1 mM methoxy-dPEG36-maleimide and incubated at room temperature for 30 minutes. In parallel reactions, purified TMcin-G1905 was treated with 0.88 M β-mercaptoethanol, 100 mM dithiothreitol, or 10 mM TCEP without addition of methoxy-dPEG36-maleimide.

### Bioinformatic analyses

AlphaFold3 (*37*) was performed on the High-Performance Computing cluster at the University of Maryland, Division of Information Technology.

Membrane protein topologies were predicted using DTU/DeepTMHMM version 1.0.44 (*38*). Topology maps were generated using TOPO2 and modified using Adobe Illustrator.

The hmmscan webserver (https://www.ebi.ac.uk/Tools/hmmer/search/hmmscan) was used to detect TmcP protein families. The HMM database was Pfam 37.2 and the significant E-values were set to 0.01 and 0.03 for Sequence and Hit, respectively.

DUF3267-containing proteins were retrieved from InterPro/Pfam (PF11667), and all proteins with a pre-generated structural model were downloaded from the AlphaFold Protein structure database (https://alphafold.ebi.ac.uk/). The DUF3267 domains were extracted and aligned to the DUF3267 envelope of TmcP or the full-length TmcE structure using TM-align (*15*). In parallel, full-length protein sequences analyzed using DTU/DeepTMHMM to predict the number of transmembrane helices.

## Acknowledgements

Research reported in this publication was supported by the National Institute Of Allergy And Infectious Diseases of the National Institutes of Health under Award Number R21AI178268. The content is solely the responsibility of the authors and does not necessarily represent the official views of the National Institutes of Health. This work was also supported by University of Maryland startup funds (to S.W.D.).

## Author Contributions Statement

**Fauzia H. Nur:** Methodology, Investigation, Data Curation, and Writing – Review & Editing. **Ama N. Antwi:** Methodology, Investigation, and Data Curation. **Seth W. Dickey:** Conceptualization, Methodology, Investigation, Supervision, Project administration, Visualization, Writing – Original Draft, Writing – Review & Editing, and Funding acquisition.

