## Supplemental figures and tables for "TMcin RiPP biosynthesis in the cellular membrane"

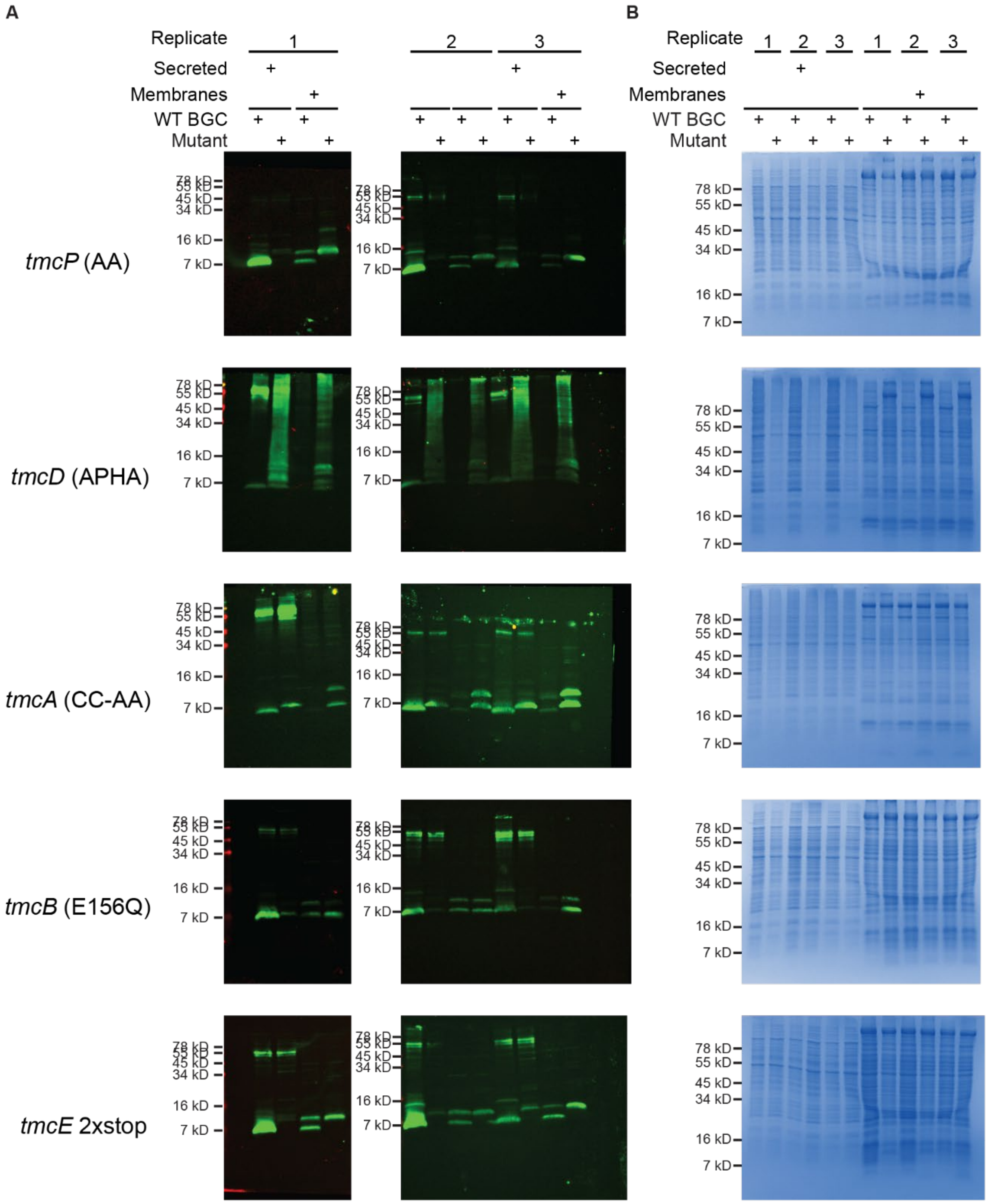


**Fig. S1: Replicates of G1905 $P_{tet}$-TMcinBiosynthesis cells after induction of TMcin biosynthesis.** (*A*) Western blots using αTMcin-G1905 and (*B*) Coomassie-stained SDS-PAGE gels of subcellular fractions after induction of TMcin biosynthesis (n = 3 biological replicates). SDS-PAGE gels stained with Coomassie blue.

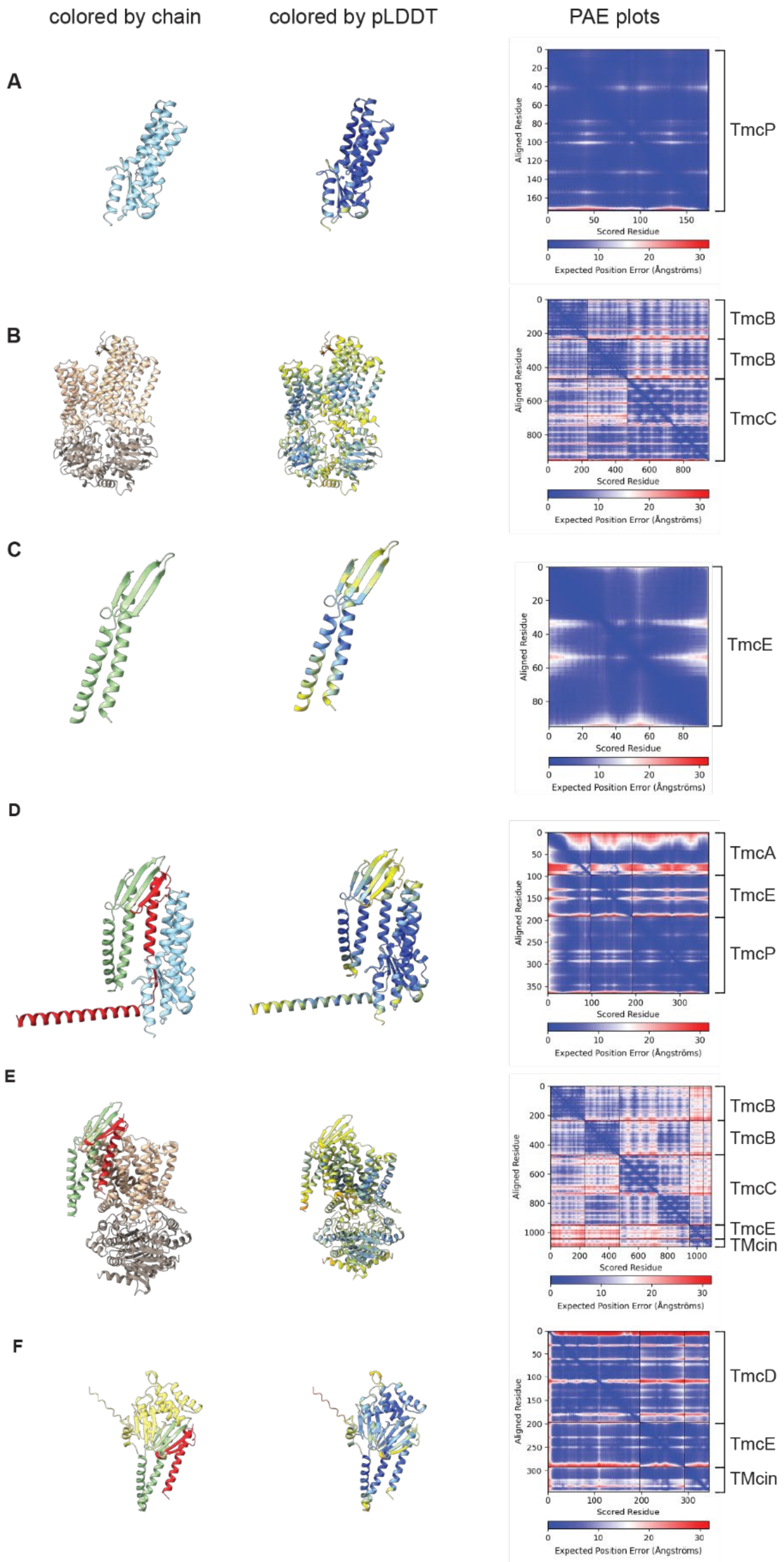


**Fig. S2**: **Confidence metrics and structures of AlphaFold3 generated models in this study.**

(*A-F*) Cartoon representation of proteins and complexes colored by chain (left) and pLDDT score (middle). Predicted aligned error plots are on the right. (*A*) TmcP (one copy, sky blue). (*B*) TmcBC with two copies of TmcB (brown) and one copy of TmcC (light brown). (*C*) TmcE (one copy, sage green), (*D*) TmcE-TmcA-TmcP complex. One copy each of TmcE (light green), TmcA (red), and TmcP (cyan). (*E*) TmcE-TMcin-TmcBC complex. (*F*) TmcD-TmcE-TMcin complex. One copy each of TmcD with the predicted signal peptide removed (yellow, residues 21-217), TMcin (red), and TmcE (light green).

**Table S2: Results from HMMScan using the TmcP sequence as input.**

| Family id | Family Accession | Clan | Env. Start | Env. End | Model Start | Model End | Bit Score | Ind. E-value | Cond. E-value | Description |
|---|---|---|---|---|---|---|---|---|---|---|
| DUF3267 | PF11667 | CL0126 | 61 | 171 | 2 | 100 | 26.5 | 8.50e-06 | 1.06e-09 | Putative zincin peptidase |

**Table S3: Oligonucleotides used in this study.**

| Oligonucleotide name | Sequence (5’-3’) | Purpose |
|---|---|---|
| pCT545_antitoxin_forward | CATATGAGCACAAACCCAAA | G1905 ΔpCT545 |
| pCT545_2Stop_antitoxin_reverse | TAATACTTAATATTTGTTTTGATAATAGCACCG | G1905 ΔpCT545 |
| 1kb_*tetML*_Forward | AATCGAATGGAAAGTCTAAAAGG | G1905 Δ*tetML* |
| *tetML*_1kb_Reverse | TTGCAGTTTTCCATCACG | G1905 Δ*tetML* |
| delta_*tetL*_Right | TAAATCGTTAAGGGATCAACTTTG | G1905 Δ*tetML* |
| delta-*tetM*_Left | GTGATTTTCCTCCATTCAAAAAC | G1905 Δ*tetML* |
| pTX15 MCS Left | GATATCCTCCTCAGCTGGATCCTACATTTTAGTTGGTTAATTT | pKX17 (introduce multiple cloning site) |
| pTX15 MCS Right | ATATGATTTGCGGCCGCATACGCGTGTGATGAACGCTTTATCTTAATGC | pKX17 (introduce multiple cloning site) |
| pKX17_delta_PxylA_ApaI_Right | TATGCTGGGCCCAGCTGAGGAGGATATCATAT | pKT17 (remove *xylR* and $P_{xylA}$) |
| pKX17_delta_*xylR*_Left | GATCATCATGACAGATCCGG | pKT17 (remove *xylR* and $P_{xylA}$) |
| pIMAY_*tetR*_Reverse | TGAAGTTACCATCACGGA | pKT17 (*tetR* and $P_{tet}$) |
| pIMAY_Pxyl_ApaI_Forward | CTGAGGGCCCCTCTATCAATGATAGAGAGCTTATTTTAATTATACTCTATCAATGATAG | pKT17 (*tetR* and $P_{tet}$) |
| *tetR*_remove_NdeI_Left | TGCGGATTAGAAAAACAACTTAAAT | pKT17 (remove NdeI site in *tetR*) |
| *tetR*_remove_NdeI_Right | aattATCAATTCAAGGCCGAATAAG | pKT17 (remove NdeI site in *tetR*) |
| *tmcB*_NdeI_Forward | TCATGTCATATGTTGAAAATTGAAAATGTATCATTTAAGTATAC | pKT17 BGC-core: insert *tmcB-tmcA* |
| *tmcA*_NotI_Reverse | ATACCAGGCGGCCGCTTATCTTTTGCACTCAACATCG | pKT17 BGC-core: insert *tmcB-tmcA* |
| pRB_Δ_GramN_Left | GCATCTGTGCGGTATTTCA | Delete Gram-negative replication and antibiotic resistance marker |
| pRB_Δ_GramN_Right | GTCATTACCCCAGGCGTTTA | Delete Gram-negative replication and antibiotic resistance marker |
| Δ_*tmcA*_Right | TAACTTTTAAACTACAAATGCGTGG | Delete *tmcB-tmcA* in pRB5gp + BGC |
| Δ_*tmcB*_Left | TTGTAAGTCGTCCTTTAATTTTTTTAAATATATATAATATTG | Delete *tmcB-tmcA* in pRB5gp + BGC |
| *tmcD*_APHA_left | TGCGTGAGGAGCTTTAAAATCTTCGAAAACTATAATTTTATTTTTTGAAC | *tmcD* APHA mutagenesis |
| *tmcD*_R70_Right | AGAACTCTATATAATAATATTATAAAAAAAATTGATGATGG | *tmcD* APHA mutagenesis |
| *tmcB*_E156Q_Forward | CAAACATTAAATGGAATGGATATTGAATCA | *tmcB* E156Q mutagenesis |
| *tmcB*_D155_Reverse | ATCTATTATTAGAATATTGGGTGCTATTAAC | *tmcB* E156Q mutagenesis |
| *tmcE*_S10L11Stop_Forward | TAATAGGTTTTTTCTATAGTAACTACCGGAAT | *tmcE* 2xstop mutation |
| *tmcE*_N9_Rev | GTTAATTAAATATATTAAGTGTTTCATTTCTCGAC | *tmcE* 2xstop mutation |
| *tmcP*_H68A_Forward | GCAGAATATATGCATATATATTTCTTGAAGAAAAAGC | *tmcP* H68A mutagenesis |
| *tmcP*_H68A_Reverse | TATAGCAAAAGAAAGAAGCAAACC | *tmcP* H68A mutagenesis |
| *tmcP*_D158A_Forward | GCAGGAGTAATGTTAATCAAAGGAATATTAaag | *tmcP* D158A mutagenesis |
| *tmcP*_D158A_Reverse | TCCTAATGGTGGAAGTATGTTAA | *tmcP* D158A mutagenesis |
| *tmcA*_C70A_Right | GCAGTAGTGAATCAACATGGTAAGTT | *tmcA* C70A mutagenesis |
| *tmcA*_C70A_Left | CCAAATAGTTAAACCAGCAAAAATT | *tmcA* C70A mutagenesis |
| *tmcA*_C95A_Left | TGCCTCAACATCGAGGCTAAC | *tmcA* C95A mutagenesis |
| *tmcA*_C95A_Right | AAAAGATAAGCGGCCGCA | *tmcA* C95A mutagenesis |

**Table S4: Plasmids used in this study.**

| Plasmid Name | Source |
|---|---|
| pKX15 | (*35*) |
| pKX17 | This study |
| pKT17 | This study |
| pKT17+BGC-core | This study |
| pKT17+BGC-core *tmcB*(E156Q) | This study |
| pKT17+BGC-core *tmcE*(2xstop) | This study |
| pKT17+BGC-core *tmcP*(H68A,D158A) | This study |
| pKT17+BGC-core *tmcA*(C70A,C95A) | This study |
| pKT17+BGC-core *tmcE*(2xstop) *tmcA*(C70A,C95A) | This study |
| pKT17+BGC-core *tmcB*(E156Q) *tmcA*(C70A,C95A) | This study |
| pRB5gp | This study |
| pRB+BGC | (*6*) |
| pRB5gp+BGC | This study |
| pRB5gp+BGC-Δcore | This study |
| pRB5gp+BGC-Δcore *tmcD*(C66A,C69A) | This study |
